# Human Saphenous Vein Ex Vivo Culture as a Translational Model of Intimal Hyperplasia

**DOI:** 10.1101/2025.06.17.660253

**Authors:** Shuang Zhao, Celine Deslarzes-Dubuis, Severine Urfer, Jonathan Thevenet, Clémence Bechelli, Martine Lambelet, Florent Allagnat, Sébastien Déglise

## Abstract

**BACKGROUND:** Intimal hyperplasia (IH) significantly limits the long-term patency of saphenous vein grafts following bypass surgery, with no human models available to fully understand its complex pathogenesis. Although animal models, primarily murine systems, have provided mechanistic insights into IH, limitations persist in translating these findings to human pathophysiology. Here, we evaluate the translational value of a static *ex vivo* culture model using human saphenous vein segments to study IH.

**METHODS:** Human saphenous vein segments obtained from patients who underwent lower limb bypass surgery were cultured *ex vivo* for 7 days under static conditions. Histological and immunohistochemical analyses were conducted to evaluate endothelial dysfunction, vascular smooth muscle cell (VSMC) phenotype switching, extracellular matrix (ECM) remodeling, inflammation, and apoptosis. Spatial transcriptomics (GeoMx) were employed to characterize the localized transcriptional alterations, which were subsequently validated using targeted qPCR, western blotting, and additional immunostaining techniques.

**RESULTS:** Cultured vein segments developed characteristic features of IH, including marked endothelial dysfunction, increased apoptosis and proliferation, ECM remodeling and neointima formation. Spatial transcriptomics revealed localized VSMC dedifferentiation and activation of inflammatory, oxidative stress, and ECM remodeling pathways. Importantly, we also observed evidence of osteochondrogenic differentiation of human VSMCs during IH, with significant upregulation of osteogenic markers such as RunX2.

**CONCLUSIONS:** Our *ex vivo* human saphenous vein model captures the complex molecular and cellular dynamics of IH, offering insights into endothelial dysfunction, VSMC plasticity, and osteochondrogenic transitions. This translational model holds significant promise for evaluating novel therapeutic strategies targeting graft IH.

## Introduction

Intimal hyperplasia (IH) remains a major clinical challenge, significantly impacting the long-term patency of vascular grafts, particularly saphenous vein (SV) grafts used in bypass surgery. Despite advances in surgical techniques and pharmacological therapies, approximately 30–50% of vein grafts develop IH within 1-18 months post-implantation, leading to vessel stenosis and graft failure^1–2,3^. Although animal models, primarily murine systems, have provided valuable mechanistic insights into IH, considerable limitations persist, especially in translating these findings to human pathophysiology.

The pathophysiology of IH involves a complex interplay of cellular and molecular processes initiated by vascular injury and endothelial dysfunction. Endothelial injury triggers inflammatory responses, attracting inflammatory cells and promoting cytokine release, which subsequently stimulate vascular smooth muscle cell (VSMC) proliferation, migration, and dedifferentiation from a contractile to a synthetic phenotype^4,5^. This phenotypic switch results in excessive extracellular matrix (ECM) deposition and vascular remodeling, ultimately leading to neointima formation, vessel stenosis, and compromised graft patency^6–9^.

A critical gap in the field is the lack of human models that accurately capture the complexity of IH, including endothelial dysfunction, VSMC plasticity, ECM remodeling, and inflammation. To bridge this gap, we developed a model of *ex vivo* vein perfusion and demonstrated that arterial pressure promotes IH^10–12^. Recently, we further developed a simpler *ex vivo* model of static culture of human saphenous vein segments, which was successfully used to evaluate novel therapeutic approaches^12–15^. In the present study, we further characterize this model, highlighting the morphological, cellular, and molecular changes underpinning IH formation and providing insights into vascular remodeling processes and potential therapeutic targets to improve clinical outcomes.

## Methods

### Data Availability Statement

The data that support the findings of this study are available from the corresponding author upon reasonable request. Spatial omics data from the high-throughput sequencing combined with the GeoMx Digital Spatial Profiler (DSP) have been made publicly available at NCBI Gene Expression Omnibus (GEO) under the accession number : GSE299685 (GEO Accession viewer).

**Main reagents and chemical are available in the Major resource table in the supplemental data file.**

### Human vein culture

Static vein culture was performed as recently described^13,15,16^. Briefly, segments of great saphenous vein were cut in 5 mm segments randomly distributed between conditions. One segment (D0) was immediately preserved in formalin or flash frozen in liquid nitrogen and the others were maintained in culture for 7 days in RPMI-1640 Glutamax I supplemented with 10% FBS and 1% antibiotic solution (10,000 U/mL penicillin G, 10,000 U/mL streptomycin sulphate) in cell culture incubator at 37°C, 5% CO_2_ and 21% O_2_. This medium was changed every 2 days. A central, 5 mm-thick ring was cut and fixed in formalin for morphometry after 7-day in culture. The remaining fragments were frozen and reduced into powder for RT-PCR and Western blot analysis. Eight veins were obtained from randomly selected patients who underwent lower limb bypass surgery for critical ischemia.

### Histology

After 7 days in culture, or immediately upon vein isolation (D0), human vessel segments were fixed in buffered formalin, embedded in paraffin and cut into 6 µm sections, and stained with Van Gieson Elastic Laminae (VGEL) or Polychrome Herovici as previously described^15,17^. Calcium deposits were stained using a 2 % w/w Alizarin Red Solution in water for 3 minutes before washing and dehydration in 100% acetone, followed by Acetone-Xylene (1:1) solution and final clearing in 2 times 5 min in pure xylene. Slides were scanned using a ZEISS Axioscan 7 Microscope Slide Scanner.

For intimal and medial thickness, 96 (4 measurements/photos and 4 photos per cross section on six cross sections) measurements were performed^15^. Two independent researchers blinded to the conditions did the morphometric measurements using the ImageJ 1.53t software (NIH, USA). Immunohistochemistry was performed on paraffin sections. After rehydration and antigen retrieval (TRIS-EDTA buffer, pH 9.0, 1 min in an electric pressure cooker autocuiser Instant Pot duo 60 under high pressure), immunostaining was performed on human vein sections using the rabbit-specific HRP/DAB detection IHC detection kit (ab236469) according to manufacturer’s instructions. Slides were further counterstained with hematoxylin. The positive immunostaining area was quantified using the Fiji (ImageJ 1.53t) software and normalized to the total area of the tissue by two independent observers blinded to the conditions. The antibodies used in the study are described in Supplemental **Table S2**.

### Western blotting

Vessels were flash-frozen in liquid nitrogen, grinded to powder and resuspended in SDS lysis buffer (62.5 mM TRIS pH6.8 5% SDS, 10 mM EDTA). Protein concentration was determined by DC protein assay (Biorad AG, Switzerland). 10 to 20 µg of protein were loaded per well. Lysates were resolved by SDS-PAGE and transferred to a PVDF membrane (Immobilon-P, Millipore AG, Switzerland). Immunoblot analyses were performed as previously described^16^ using the antibodies described in Supplemental **Table S2**. Membranes were revealed by enhanced chemiluminescence (Immobilon, Millipore) using the Azure 280 device (Azure Biosystems AG, Switzerland) and analyzed using Fiji (ImageJ 1.53t). Protein abundance was normalized to total protein using Pierce™ Reversible Protein Stain Kit for PVDF Membranes (Thermo Fisher Scientific AG, Switzerland).

### Reverse transcription and quantitative polymerase chain reaction (RT-qPCR)

Flash frozen vessels powder was homogenized in TripureTM Isolation Reagent (Roche, Switzerland), and total RNA was extracted according to the manufacturer’s instructions. After RNA reverse transcription (Prime Script RT reagent, Takara), cDNA levels were measured by qPCR Power-Up SYBR™ Green Master Mix (Ref: a25742, Applied Biosystems, Thermo Fischer Scientific AG, Switzerland) in a Quant Studio 5 Real-Time PCR System (Applied Biosystems, Thermo Fischer Scientific AG, Switzerland), using the primers given in the Supplemental **Table S3**.

### GeoMx Digital Spatial Profiling (DSP) Slides Preparation

FFPE tissue sections of 5-µm were prepared for RNA profiling according to Nanostring’s instructions^18,19^. FFPE tissue sections were baked at 60 °C for 1 hour. Slides were deparaffinized in 3 xylol baths of 5 min, then rehydrated in ethanol gradient from 100% EtOH, 2 baths of 5 min followed by 95% EtOH, 5 min. Slides were then washed in PBS 1X.

Antigen retrieval was done in Tris-EDTA pH 9.0 buffer at 100°C for 15 min at low pressure. Slides were first dipped into hot water 10 seconds then dipped into Tris-EDTA buffer. Cooker vent stayed open during the procedure to ensure low pressure and reach 100°C. Slides were then washed into PBS 1X, and incubated into proteinase K in PBS (1ug/ml) for 15 min at 37°C, and washed again in PBS 1X. Tissue were post-fixed in 10% neutral-buffered formalin 5 min, washed 2 times 5 min in NBF stop buffer (0.1M Tris Base, 0.1M Glycine) and finally one time in PBS 1X. The mix of Whole Transcriptomic Atlas (WTA, Nanostring, Bruker, United States) probes was dropped on each section and covered with HybriSlip Hybridization Covers. Slides were then put for hybridization overnight at 37°C in a Hyb EZ II hybridization oven (Advanced cell Diagnostics, Bio-Techne, United States). The day after, HybriSlip covers were gently removed and 25-min stringent washes were performed twice in 50% formamide and 2X SSC at 37 °C. Tissues were washed for 5 min in 2X SSC, then blocked in Buffer W (Nanostring, Bruker, United States) for 30 min at RT in a humidity chamber. Next, antibody targeting MYH11 (ab224804, 1:100) in Buffer W were applied to each section for 1 h at RT. Slides were washed four times in fresh 2X SSC for 3 min and incubated 1 h at RT with Goat anti-Rabbit AF594 antibody (ab150080, 1:1000, Abcam, United Kingdom) and 500 nM Syto13. Slides were washed twice in fresh 2X SSC then loaded on the GeoMx Digital Spatial Profiler (DSP).

For slides collection, entire slides were imaged at ×20 magnification and morphologic markers were used to select Region of Interest (ROI) using organic shapes. Automatic segmentation of ROI based on MYH11 markers were used to defined Area of Illumination (AOIs). AOIs were exposed to 385 nm light (UV), releasing the indexing oligonucleotides which were collected with a microcapillary and deposited in a 96-well plate for subsequent processing. The indexing oligonucleotides were dried down overnight and resuspended in 10 μl of DEPC-treated water.

NGS sequencing platform was used as readout. Sequencing libraries were generated by PCR from the photo-released indexing oligos and AOI-specific Illumina adapter sequences, and unique i5 and i7 sample indices were added. Each PCR reaction used 4 μl of indexing oligonucleotides, 4 μl of indexing PCR primers, 2 μl of Nanostring 5X PCR Master Mix. Thermocycling conditions were 37 °C for 30 min, 50 °C for 10 min, 95 °C for 3 min; 18 cycles of 95 °C for 15 seconds, 65 °C for 1 minutes, 68 °C for 30 seconds; and 68 °C for 5 minutes. PCR reactions were pooled and purified twice using AMPure XP beads (A63881, Beckman Coulter, United States), according to the manufacturer’s protocol. Pooled libraries are paired-sequenced at 2 × 27 base pairs and with the single-index workflow on an Illumina Novaseq instrument.

Novaseq-derived FASTQ files for each sample were compiled for each compartment using the bcl2fastq program of Illumina, and then demultiplexed and converted to digital count conversion files using the GeoMx DnD pipeline v2.0.0.16 of NanoString according to manufacturer’s pipeline.

### Quality control and Normalization of GeoMx DSP Data

QC and Q3 normalization were performed using the R package GeoMxTools v3.4.0. Probes were checked for outlier status. A probe was removed if the geometric mean of that probe’s counts from all segments divided by the geometric mean of all probe counts representing the target from all segments was less than 0.1 or if the probe was an outlier according to the Grubb’s test in at least 20% of segments. The counts for all remaining probes for a given target were then collapsed into a single metric by taking the geometric mean of probe counts. Genes were filtered out with abnormal low signal. Genes with less than 1% of segments detected and below the limit of quantification (LOQ) set at 2 geometric standard deviation above the geometric mean were removed from the study. Gene counts were normalized to the geometric mean of the 75^th^ percentile across all AOIs to give the upper quartile (Q3) normalization factors for each AOI.

### GeoMx digital spatial profiling (DSP) analysis

The visualization of categorical variable flow was performed using the easylluvial package v0.3.2 in R to generate a Sankey diagram. Uniform Manifold Approximation and Projection (UMAP) was conducted using the umap package v0.2.10.0 Differential gene expression between groups of segments was assessed on Q3 normalized data using linear mixed-effects models (mixedModelDE). Enrichment mapping and gene set enrichment analysis (GSEA) using Hallmark gene sets from the Molecular Signatures Database (MSigDB, collection H) were performed with ClusterProfiler v4.8.1. To estimate cell abundance, cell type deconvolution from gene expression data was performed using SpatialDecon with an adapted version of DeMicheli Human Muscle Atlas cell profile matrix. Ligand-to-target analysis were conducted via the NicheNet package v2.2.0 ^20^.

### Statistical analyses

All experiments adhered to the ARRIVE guidelines and followed strict randomization. All experiments and data analysis were conducted in a blind manner using coded tags rather than actual group name. All experiments were analyzed using GraphPad Prism 9 or R. Normal distribution of the data was assessed using Kolmogorov-Smirnov tests. All data had a normal distribution. Unpaired bilateral Student’s t-tests, Wilcoxon test, or one way ANOVA were performed followed by multiple comparisons using post-hoc t-tests with the appropriate correction for multiple comparisons.

### Ethics Statement

Human great saphenous veins were obtained from donors who underwent lower limb bypass surgery^21^. Written, informed consent was obtained from all vein donors for human vein cultures. The study protocols for organ collection and use were reviewed and approved by the Centre Hospitalier Universitaire Vaudois (CHUV) and the Cantonal Human Research Ethics Committee (http://www.cer-vd.ch/, no IRB number, Protocol Number 170/02), and are in accordance with the principles outlined in the Declaration of Helsinki of 1975, as revised in 1983 for the use of human tissues.

## Results

### Human saphenous vein segments develop intimal hyperplasia after static ex vivo culture

Static culture of human vein segments for 7 days (D7) led to prominent morphological alterations indicative of intimal hyperplasia (IH). Histological staining with VGEL and Herovici revealed increased intimal thickness and enhanced extracellular matrix (ECM) deposition compared to freshly isolated control segments (D0) (**Fig. 1A-B**).

**Figure 1.**
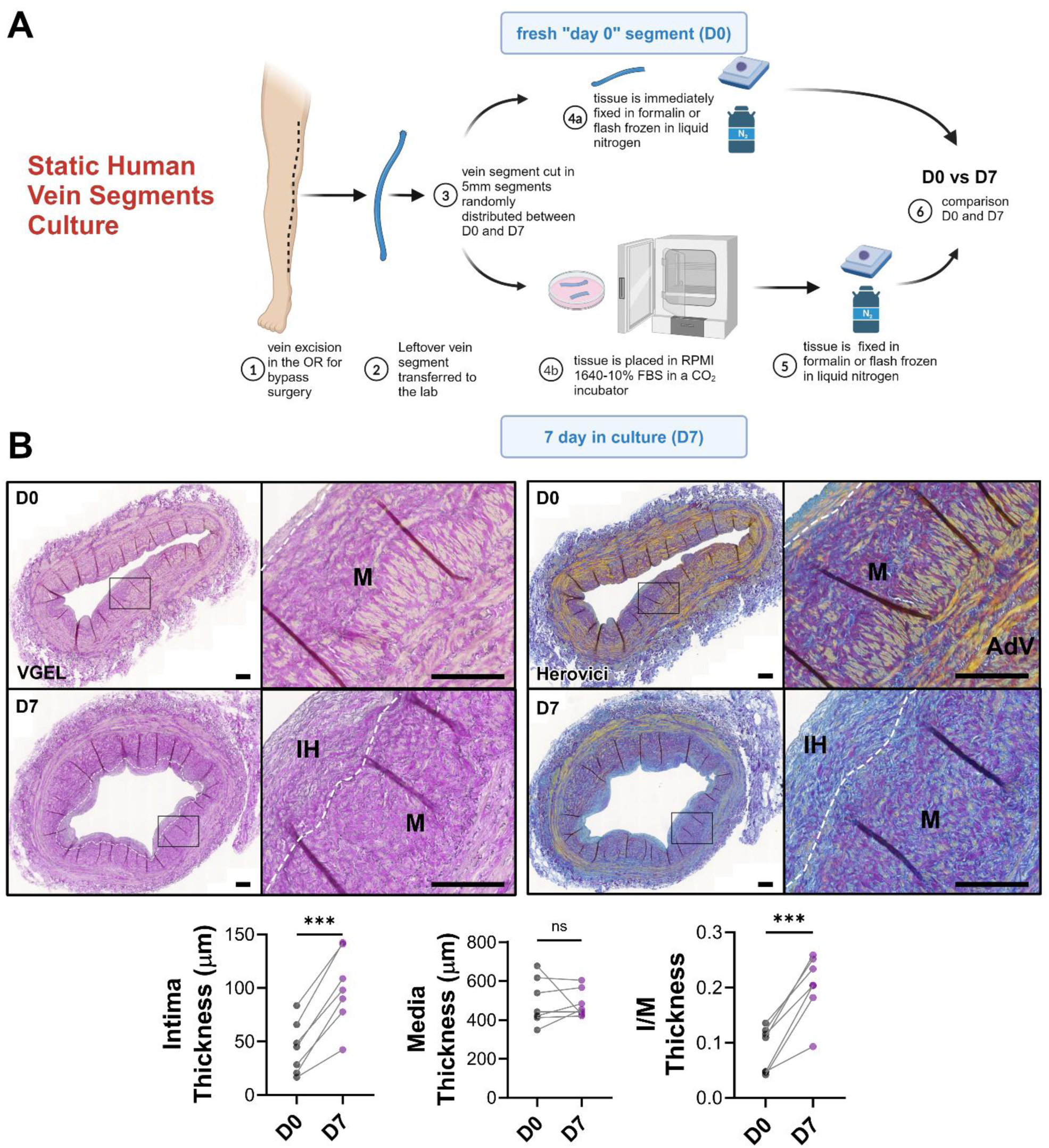
Human vessel segments develop intimal hyperplasia after 7 days in culture. (A) Schematic outline of an *ex vivo* model of intimal hyperplasia in human vessel segments. (B) Representative images of histological VGEL (left) and Herovici staining (right) and morphometric measurements of intima thickness, media thickness, intima over media ratio (I/M) of freshly isolated human vein segments (D0) or after 7 days (D7) in static culture. 5-8 different veins/patients. * p < 0.05, ** p < 0.01, *** p < 0.001, as determined by bilateral paired t-test. IH: intima hyperplasia; M: media; AdV: adventitia.

### Endothelial dysfunction and phenotypic switching of vascular smooth muscle cells (VSMCs)

As endothelial dysfunction is a key initiator of IH, we first assessed endothelial cell (EC) status. Expression of endothelial nitric oxide synthase (eNOS), critical for endothelial function, was significantly reduced at both mRNA and protein levels after 7 days in culture, despite preservation of endothelial cell lining (von Willebrand factor, VWF staining), indicating dysfunction rather than cell loss (**Fig. 2A-C**). In a healthy vessel, medial VSMC express an array of contractile markers (e.g. ACTA2, MYH11, SM22α and calponin)^22^, which are reduced upon vascular injuries and VSMC dedifferentiation^23^. Here, VSMCs underwent significant phenotypic switching, characterized by decreased expression of contractile markers including mRNA expression of ACTA2, MYH11, Calponin, and SM22α at (**Fig. 2D**). Immunohistochemical analysis confirmed diminished contractile phenotype (**Fig. 2E-J**), consistent with VSMC dedifferentiation into a synthetic phenotype associated with IH.

**Figure 2.**
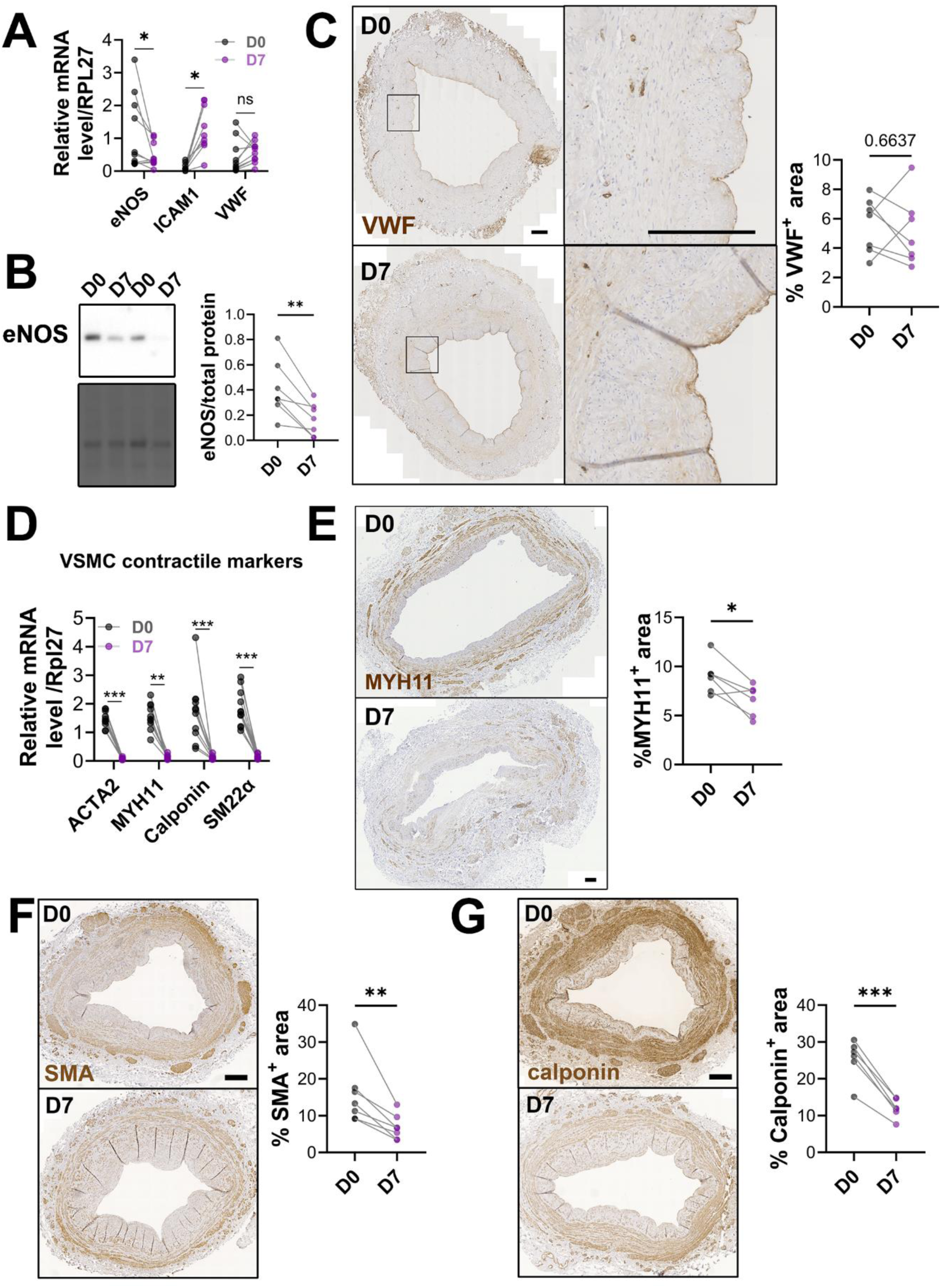
Human vessel segments exhibit EC dysfunction and a loss of VSMC contractile properties after 7 days in culture. (A) Relative mRNA expression of eNOS, ICAM1 and VWF to Rpl27 in the human vein segments at D0 and D7. (B) Representative western blot of phosphorylated eNOS and quantitative assessment of human veins at D0 and D7. (C) Representative images and quantifications of VWF expression in D0 and D7 human vein segments. Scale bars represent 200µm. (D) Relative mRNA expression of ACTA2, MYH11, Calponin and SM22α to Rpl27 in the human vein segments at D0 and D7. (E) Representative images and quantifications of MYH11 expression in D0 and D7 human vein segments. Scale bar represent 200µm. (F) Representative images and quantifications of SMA expression in D0 and D7 human vein segments. Scale bar represent 500µm. (G) Representative images and quantifications of Calponin expression as assessed by immunohistochemistry in D0 and D7 human vein segments. Scale bar represent 500µm. 5-8 different veins/patients. * p < 0.05, ** p < 0.01, *** p < 0.001, as determined by bilateral paired t-test.

### Spatial transcriptomics identifies transcriptional remodeling during IH formation

To investigate transcriptional changes underpinning IH, GeoMx spatial transcriptomics were performed, comparing D0 and D7 vein segments. The spatial analysis pipeline, including sample mounting, imaging, and data analysis, was conducted using the NanoString platform. Then we selected regions of interest (ROIs) based on MYH11^+^ staining and histopathologic architecture consistent with human vein tissue (**Fig. 3A**; **Fig. S1A**).

**Figure 3.**
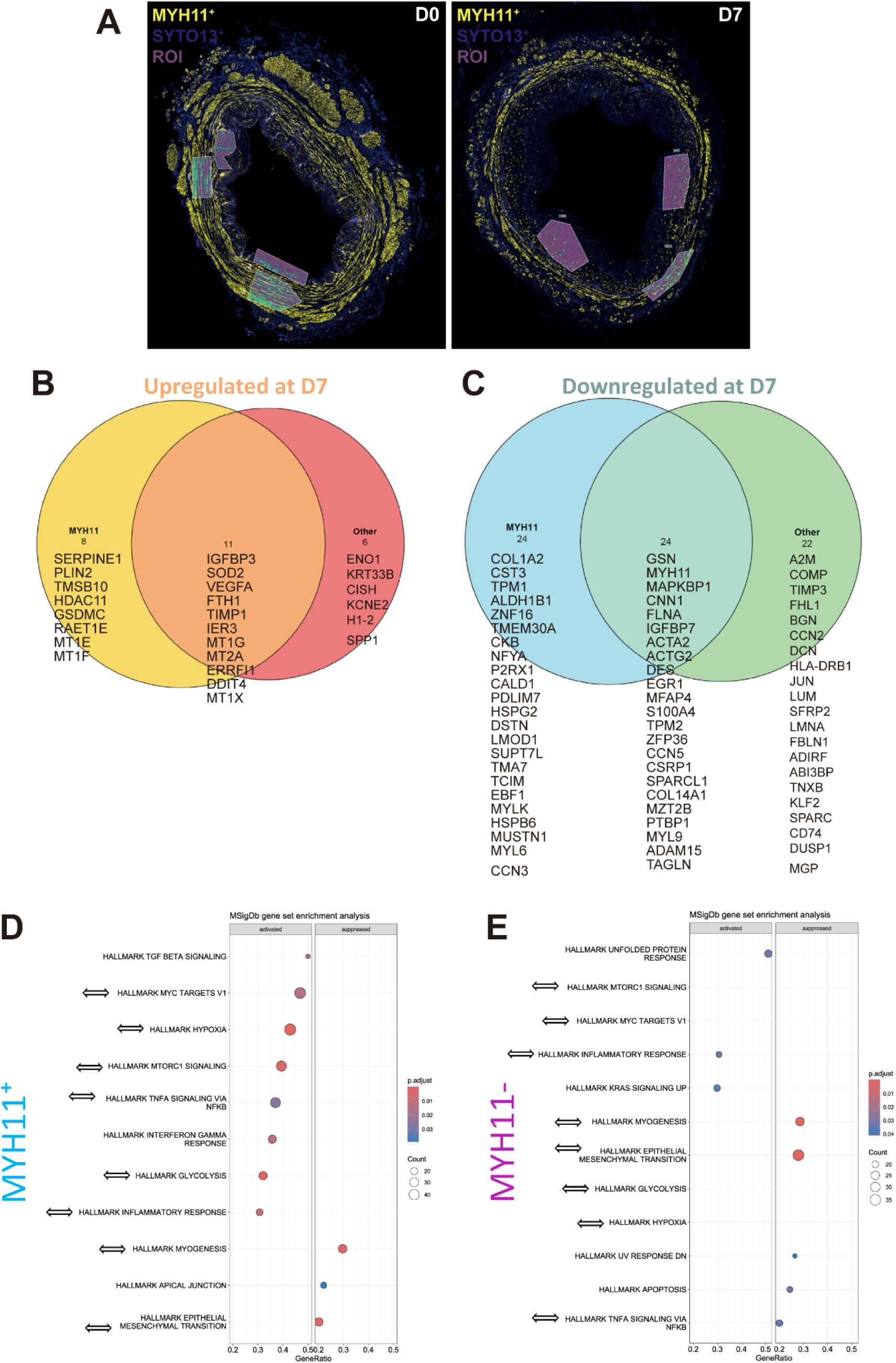
Spatial transcriptomics reveals unique pathological transcriptional differences in the human vein segments. (A) Representative immunostaining images of human vein segments at D0 and D7. VSMC are labeled using the VSMC marker MYH11 (yellow), while cell nuclei are stained with SYTO13 nucleic acid dye (blue). ROIs for spatial transcriptomics analysis are highlighted in purple shapes, showcasing selectively sampled areas. (B-C) Venn diagram of upregulated genes (B) and downregulated genes (C) in DEGs (differentially expressed genes) between two groups: MYH11^+^ cells and MYH11^-^ other cells of human vein segments at D7 vs. D0. (D-E) MSigDb gene set enrichment analysis of MYH11^+^ cells (D) and MYH11^-^ other cells (E) in human vein segments between at D0 and D7. Double-headed arrows highlight common pathways. Adjusted P value < 0.05.

Spatial analysis revealed major shifts in gene expression within both MYH11-positive (contractile VSMCs) and MYH11-negative (dedifferentiated VSMCs/fibroblasts) regions, highlighting extensive cellular remodeling (**Fig. 3B-C**; **Fig. S1B-D**). The main and most striking result was the overlap between MYH11^+^ and MYH11^-^ regions (**Fig. 3B-C**), with about 50% overlap in differentially expressed genes (DEG) analysis between D0 and D7. This result strongly suggested that MYH11^-^ regions were VSMC-rich regions that already lost the differentiation marker MYH11, but still express ACTA2 and CNN1 markers. The differential expression analysis revealed strong downregulation of contractile genes (MYH11, ACTA2, CNN1) accompanied by significant upregulation of genes related to oxidative stress (SOD2, MT genes), ECM remodeling (BGN, LUM, TIMP1), inflammation (VEGFA, SPP1), and stress responses (DDIT4, IER3) (**Fig. 3B-C**). Pathway enrichment analysis confirmed activation of TGF-beta, mTORC1, hypoxia responses, and TNFα-NFκB signaling, indicative of a synthetic, pro-inflammatory, and proliferative environment (**Fig. 3D-E**). Again, there was a large overlap between the enriched pathways between MYH11^+^ and MYH11^−^ cells. Further cell deconvolution analyses indicated that at Day 0, the SV was primarily composed of contractile VSMCs and fibroblasts, and synthetic VSMCs, suggesting that the initial processes of neointima formation was already underway at the time of SV implantation (**Fig. 4A-B**). At Day 7, the media and intima were still predominantly composed of fibroblasts and VSMCs and both MYH11^+^ and MYH11^-^ cells underwent a phenotypic shift toward intermediate multipotent VSMCs and pro-inflammatory cells (**Fig. 4C**). The fibroblasts at Day 0, characterized by high expression of genes related to ECM secretion (FBLN1, DCN, DUSP1), were replaced by pro-inflammatory fibroblasts at Day 7, expressing genes such as IER3 and TIMP1. Further ligand-to-target interaction analysis revealed intricate crosstalk between MYH11^+^ and MYH11^-^ cells at baseline (D0), predominantly involving ligands associated ECM remodeling (MMP9, ITGB2, VCAM1) and pro-inflammatory signaling (CXCL8), suggesting active ECM remodeling in veins from “healthy” old patients (**Fig. 4D**). By day 7 (D7), these interactions diminished significantly, suggesting disrupted cellular communication, accompanied by a shift toward heightened inflammatory signaling and enhanced ECM remodeling (**Fig. 4D**). Collectively, spatial transcriptomics highlighted distinct transcriptional profiles and enriched pathways within both cellular regions at D7, emphasizing hypoxia, inflammation, and ECM remodeling as central drivers of vascular remodeling and disease progression.

**Figure 4.**
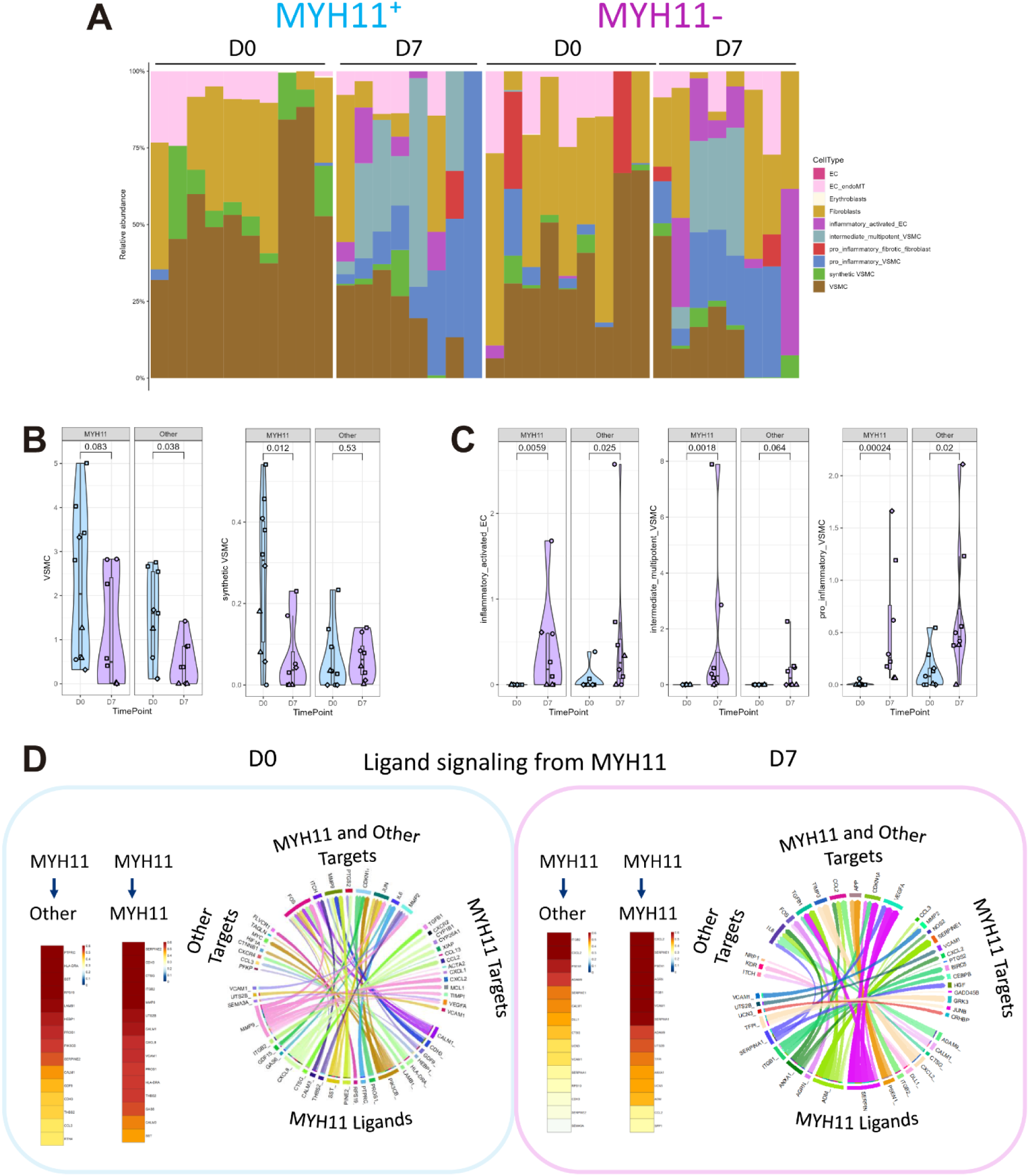
Spatial transcriptomics characterizes the transformation of VSMC and other cell types and the active ligand-signalling networks during IH progression in human vessel segments. (A) Cell type proportions grouped by D0 and D7 in MYH11^+^ and MYH11^-^ regions, respectively. (B) Violin plots of cell type proportion of VSMC (left) and synthetic VSMC (right) between D0 and D7 in the MYH11^+^ and other regions. Wilcoxon test was used to assess the significance. (C) Violin plots of cell type proportion of inflammatory activated EC (left), intermediate multipotent VSMC (middle) and pro-inflammatory VSMC (right) between D0 and D7 in the MYH11^+^ and other regions. Wilcoxon test was used to assess the significance. (D) Ligand-receptor signaling pathways from MYH11^+^ cells to MYH11^+^ and other target populations at D0 (left, blue) and D7 (right, pink). Heatmaps (left of each panel) of top ligands based on ligand activity signalling from MYH11^+^ cells to MYH11^+^ and other target populations. Circos plot (right of each panel) showing ligand signalling from MYH11^+^ ligands to MYH11^+^ and other targets.

### Oxidative stress, ER stress, apoptosis, and proliferation in VSMCs during IH development

Guided by the spatial transcriptomic findings, targeted analyses confirmed increased oxidative stress (HO-1, Nrf2, 4HNE staining) and ER stress (XBP1, CHOP, ATF4) at D7 (**Fig. 5A-C**). These stress responses coincided with significant apoptosis induction, as indicated by elevated pro-apoptotic BAX expression and reduced anti-apoptotic BCL2 (**Fig. 5D**), and increased TUNEL-positive staining (**Fig. 5E**). Despite the elevated apoptosis, PCNA staining revealed increased proliferation rates in VSMCs (**Fig. 5F**) suggesting a dynamic interplay between cell death and proliferation during IH formation.

**Figure 5.**
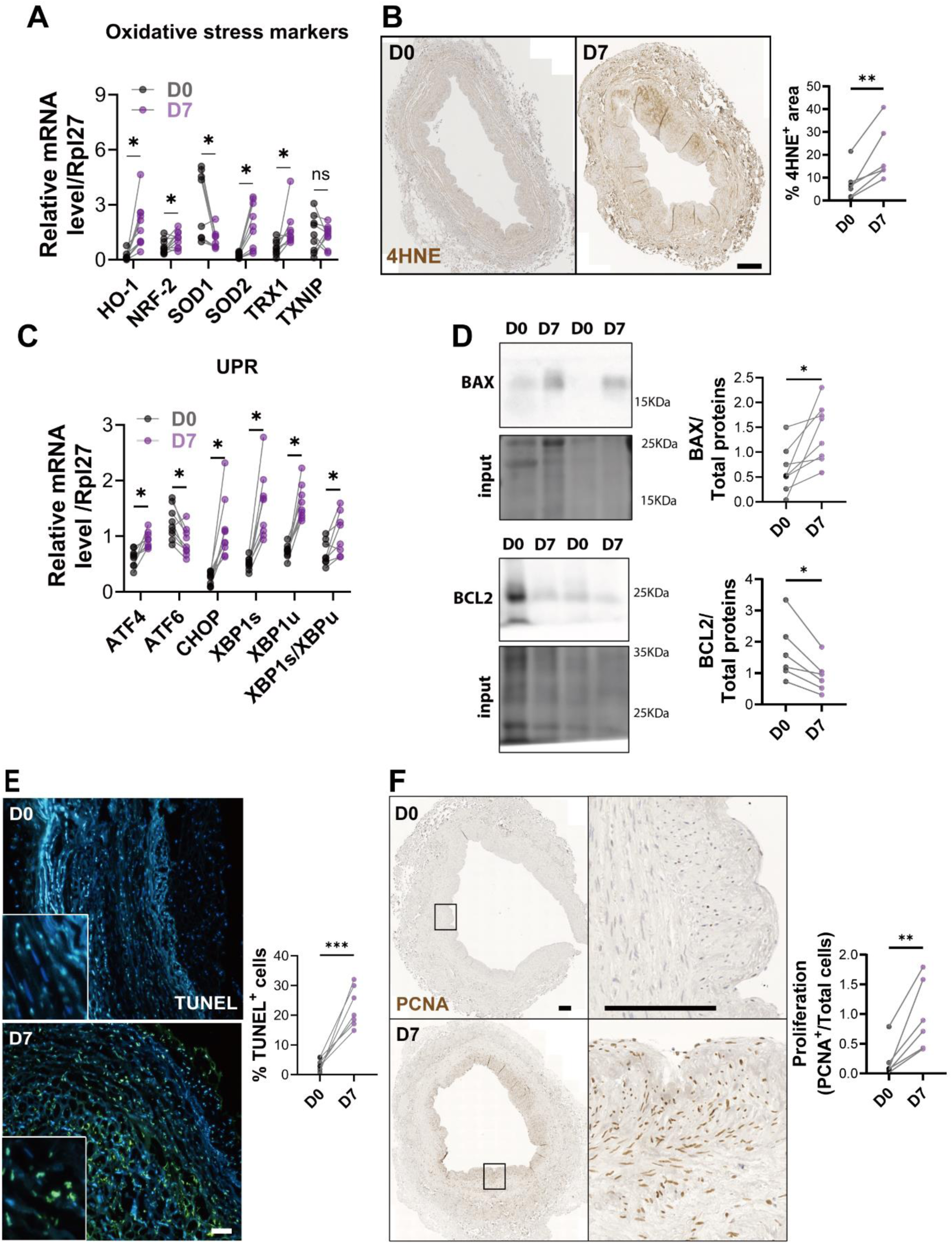
Oxidative stress, VSMC apoptosis and proliferation contribute to the progression of IH. (A) Relative mRNA expression of HO-1, NRF-2, SOD1, SOD2, TRX1 and TXNIP to Rpl27 in the human vein segments at D0 and D7. (B) Representative images and quantifications of 4HNE expression in the human vein segments at D0 and D7. Scale bar represent 500 µm. (C) Relative mRNA expression of ATF4, ATF6, CHOP, XBP1s, XBP1u, and XBP1s/XBP1u to Rpl27 in the human vein segments at D0 and D7. (D) Representative western blot of BAX, BCL2 and quantitative assessments of D0 and D7 human vein segments. (E) Representative images and quantifications of TUNEL^+^ cells. Scale bar= 50 µm. Insets are 4-fold magnification. (F) Representative images and quantifications of PCNA expression at D0 and D7. Scale bars represent 250 µm. 5-8 different veins/patients. * p < 0.05, ** p < 0.01, *** p < 0.001, as determined by bilateral paired t-test. UPR: unfolded protein response.

### Epigenetic regulation and VSMC phenotypic plasticity during IH progression

Spatial analysis suggested VSMC phenotypic switching, prompting further targeted investigation. Kruppel-like factor 4 (KLF4), essential for VSMC dedifferentiation, displayed decreased mRNA expression at D7 (**Fig. 6A**), but immunostaining indicated a shift from cytosolic to nuclear staining at D7, indicative of transcriptional activation (**Fig. 6B**). Nuclear localization of Yes-associated protein (YAP), a mediator of the synthetic VSMC phenotype, was also markedly increased (**Fig. 6C**). Enhanced expression of 5-hydroxymethylcytosine (5hmC), an epigenetic marker, further confirmed transcriptional reprogramming associated with IH (**Fig. 6D**).

**Figure 6.**
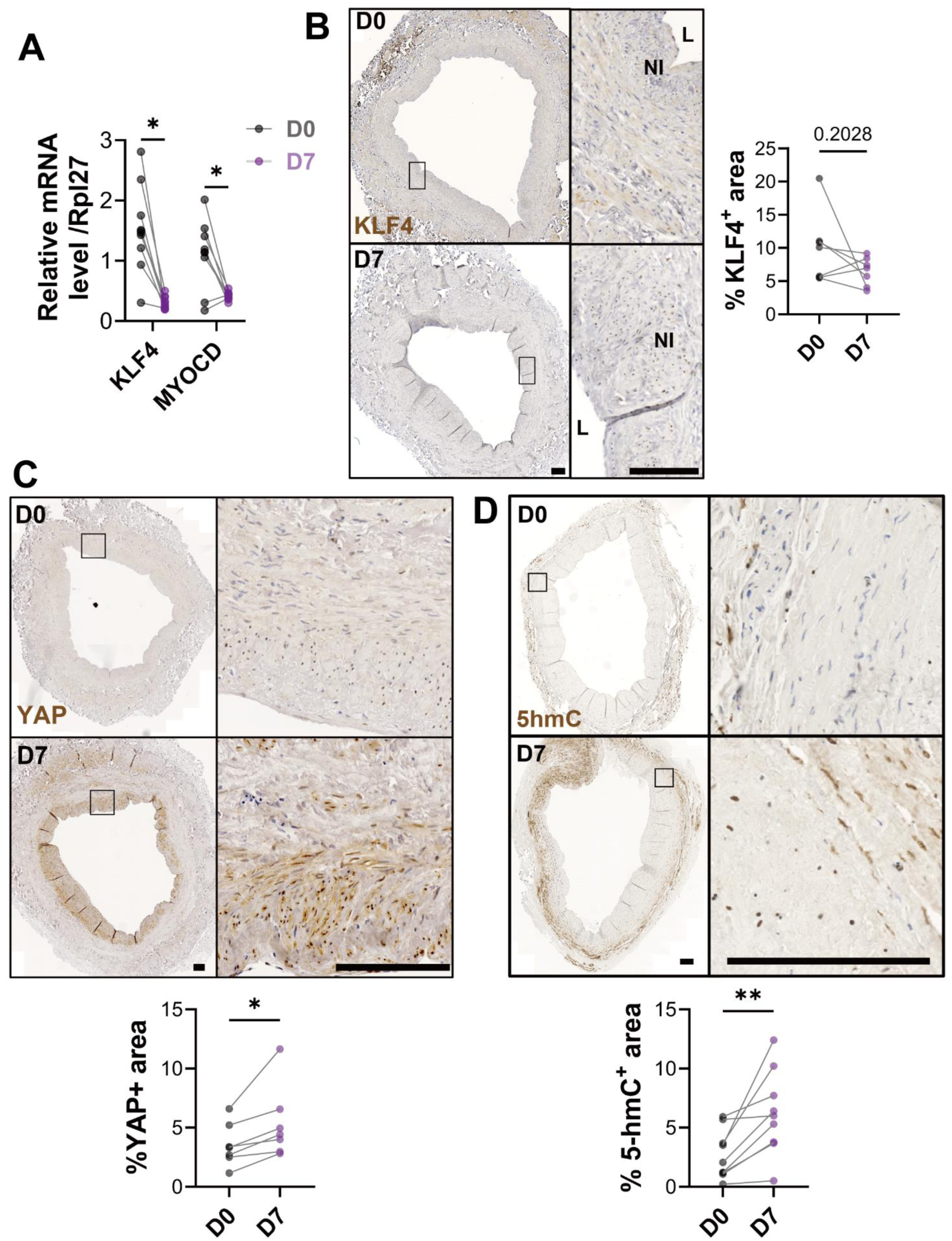
VSMC reprogramming plays a pivotal role in the development of IH. (A) Relative mRNA expression of KLF4 and MYOCD to RPL27 in the human vein segments at D0 and D7. (B-D) Representative images and quantifications of KLF4 (B), YAP (C) and 5hmC (D) expression in human veins at D0 and D7. Scale bars represent 200 µm. Data are mean ± SEM of 5- 8 different veins/patients. * p < 0.05, ** p < 0.01, as determined by bilateral paired t-test. L: lumen; NI: neointima.

### Transition of VSMCs toward a pro-inflammatory pro-fibrotic phenotype

Spatial transcriptomics suggested a shift toward ECM remodeling and pronounced inflammatory signature. Markers and activity of ECM remodeling such as proteases MMP2, MMP9, Cathepsin K were significantly elevated at D7 (**Fig. 7A-E**). Consistent with a myofibroblast-like phenotype, the fibroblast-associated HSP47 protein was increased (**Fig. 7F**). RT-qPCR also confirmed upregulation of pro-inflammatory cytokines and chemokines including IL-6, IL-1α, IL-1β, CCL5, and CXCL2 at D7 (**Fig. 7G**). Interestingly, immunostaining for Galectin-3, indicative of macrophage-like inflammatory cells, did not increase significantly, and no CD68^+^ macrophages were detected (data not shown), suggesting that inflammation was predominantly mediated by activated VSMCs rather than immune cell infiltration.

**Figure 7.**
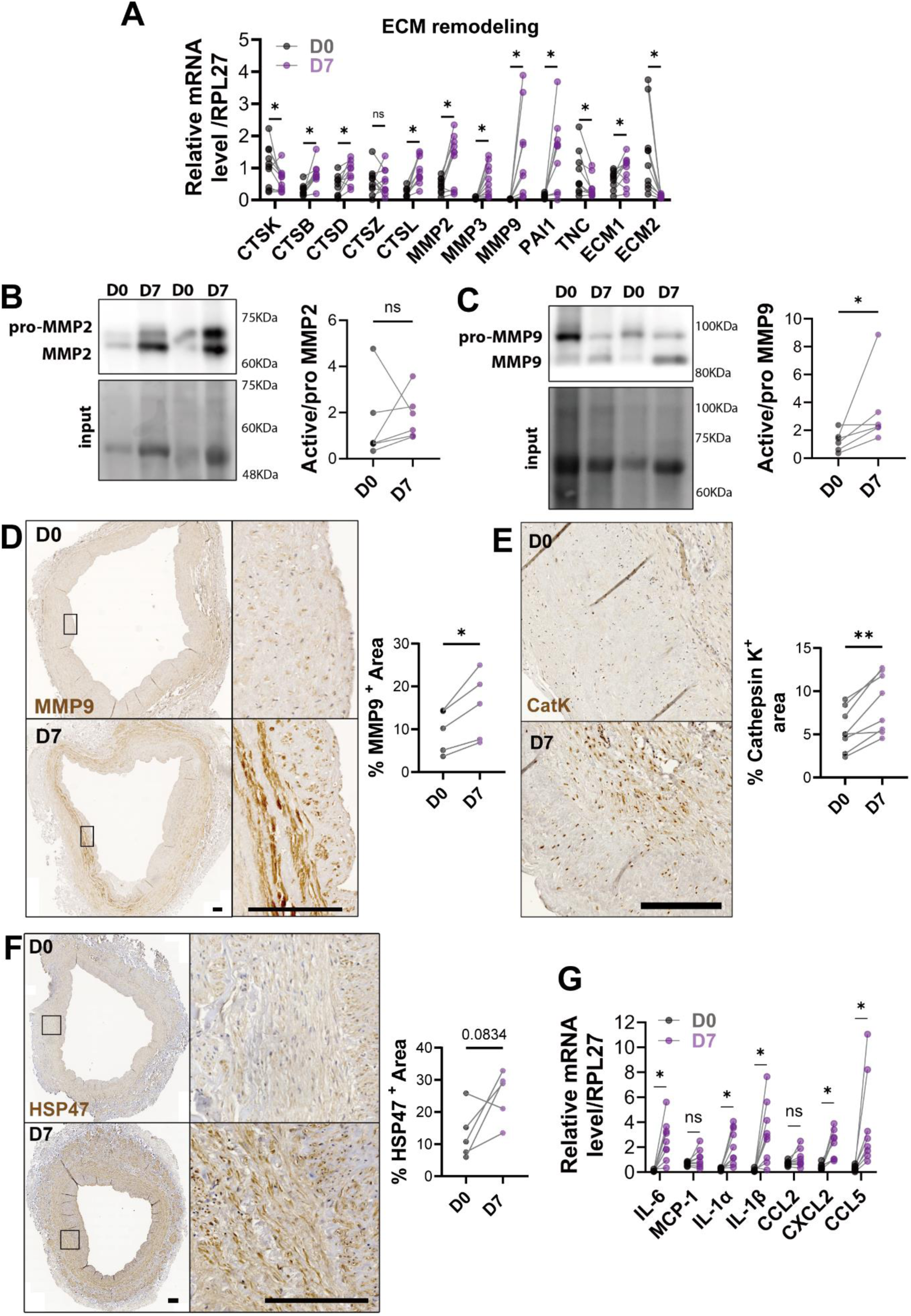
VSMC switches to a pro-inflammatory pro-fibrotic phenotype in IH. (A) Relative mRNA expression of ECM remodeling markers (MMP2, MMP3, MMP9, ECM1, ECM2, tPA, uPA, PAI1, TNC, CTSB, CTSD, CTSK, CTSL V2-8 and CTSZ) to RPL27 in the human vein segments at D0 and D7. (B-C) Representative western blot and quantitative assessments of MMP2 and MMP9 (C) expression in human veins at D0 and D7. (D-F) Representative images and quantifications of MMP9 (D), CathepsinK (E), and HSP47 (F) expression in human veins at D0 and D7. Scale bars represent 200 µm. (G) Relative mRNA expression of pro-inflammatory markers (IL-6, MCP-1, IL-1α, IL-1β, CCL2, CXCL2 and CCL5) to Rpl27 in the human vein segments at D0 and D7. 5-8 different veins/patients. * p < 0.05, ** p < 0.01, *** p < 0.001, as determined by bilateral paired t- test.

### Osteochondrogenic signature contributes to IH progression

Finally, the osteochondrogenic phenotype has been proposed to contribute to IH^24,25^. Osteochondrogenic differentiation was confirmed by increased mRNA and protein expression of key transcription factors (RunX2, Sox9) (**Fig. 8A-B**), and downstream expression of Sox9 target protein Osteopontin (**Fig. 8C**). In line with this, veins displayed signs of early calcification at D7 as assessed by Alizarin red staining (**Fig. 8D**).

**Figure 8.**
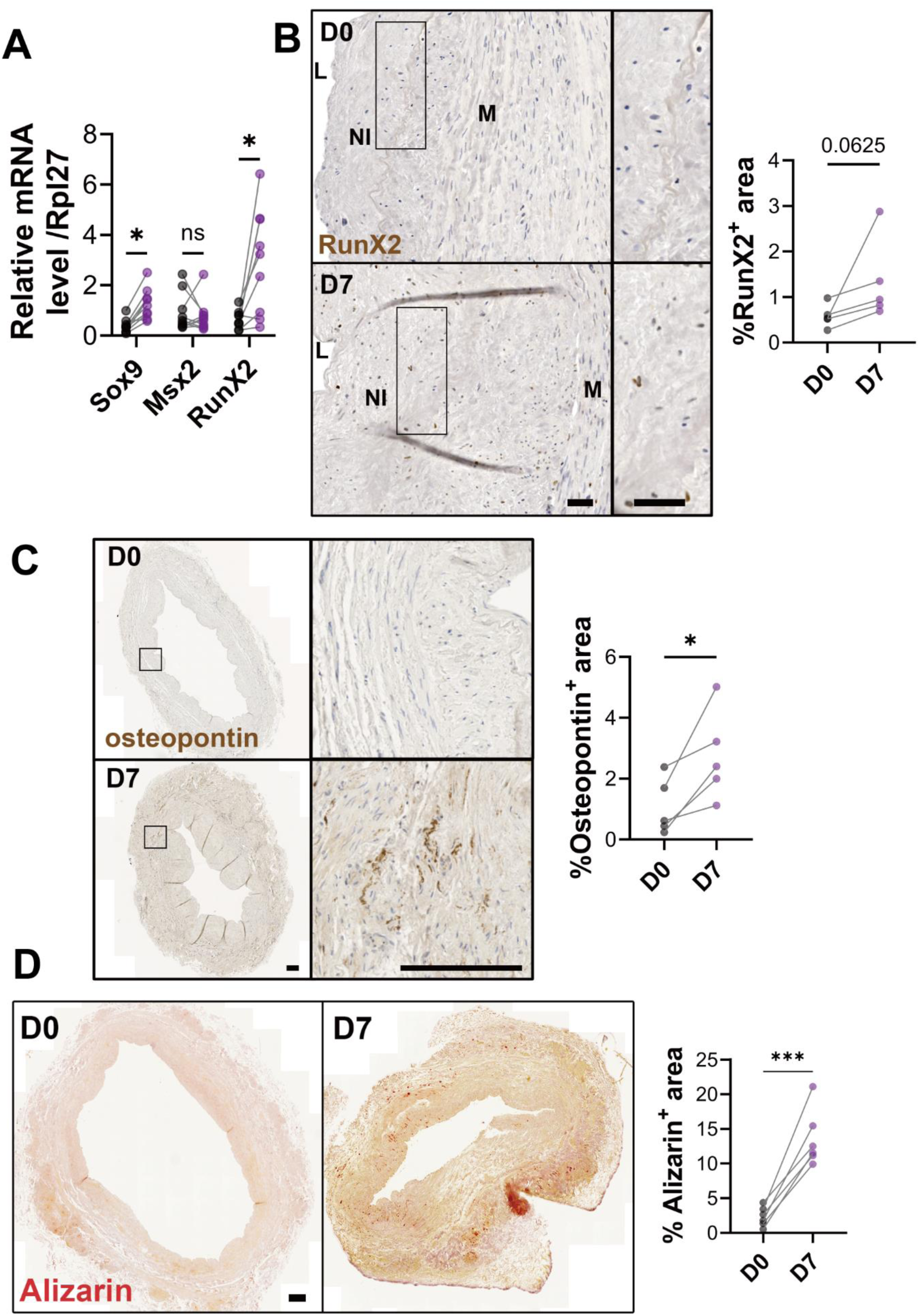
VSMC transitions to an osteochondrocyte-like phenotype in IH. (A) Relative mRNA expression of osteochondrogenic markers (Sox9, Msx2 and RunX2) to Rpl27 in the human vein segments at D0 and D7. (B-D) Representative images and quantifications of RunX2 (B), Osteopontin (C), and micro-calcification (Alizarin staining; D) in D0 and D7 human vein segments. 5-8 different veins/patients. * p < 0.05, ** p < 0.01, *** p < 0.001, as determined by bilateral paired t-test. L: Lumen; NI: neointima; M: media.

Collectively, these results illustrate a complex interplay of endothelial dysfunction, oxidative and ER stress, phenotypic plasticity, inflammation, and ECM remodeling underpinning IH formation in human vein segments *ex vivo*, validated by spatial transcriptomic insights and targeted molecular analyses.

## Discussion

Our study establishes and characterizes a robust human saphenous vein model of IH formation under *ex vivo* static culture conditions. This model recapitulates critical morphological, cellular, and molecular events relevant to human vein graft pathology. Key findings include significant endothelial dysfunction, phenotypic switching of VSMCs, oxidative and ER stress, apoptosis and proliferation dynamics, inflammation, ECM remodeling, and osteochondrogenic differentiation.

Endothelial dysfunction emerges as an early event in IH progression, evidenced by a substantial reduction in eNOS expression despite the maintained endothelial cell lining. Impaired endothelial function, notably through reduced nitric oxide (NO) bioavailability, promotes vascular inflammation and contributes to subsequent pathological remodeling^26^. EC lining of vessels require high laminar shear stress to release NO by eNOS and maintain an anti-inflammatory, anti-thrombotic phenotype^27,28^. Here, removal of blood flow likely mimics the disturbed turbulent blood flow associated with endothelial dysfunction^29^. Consistent with prior reports, our results support endothelial dysfunction as an essential initiator of IH^30^.

We observed substantial VSMC dedifferentiation characterized by decreased expression of classical contractile markers, such as ACTA2, MYH11, Calponin, and SM22α. Such phenotypic transitions from contractile to synthetic states have been widely implicated in vascular remodeling processes^4,5,7,22,31^. Supporting these observations, GeoMx spatial transcriptomics provided insights into the transcriptomic dynamics, highlighting the transcriptional remodeling underpinning VSMC dedifferentiation. Our model supports epigenetic reprogramming associated with IH, as evidenced by increased 5-hydroxymethylcytosine (5hmC) levels. This epigenetic regulation may drive VSMC transition from a contractile to a synthetic state^5^. Indeed, prior research emphasizes the significance of epigenetic regulation, including DNA methylation and hydroxymethylation, in mediating VSMC response to injury and environmental cues^32,33^. Epigenetic modifications and dedifferentiation are further supported by the identification of MSC-like cells with nuclear localization of transcriptional regulators KLF4 and YAP, and down-regulation of MYOCD. KLF4 has previously been identified as a critical regulator of VSMC phenotypic plasticity^34–37^. In animal models, this plasticity has been proposed to lead to the formation of pro-fibrotic^8^ and pro-inflammatory phenotypes^38,39^. Many myofibroblast-like VSMC are derived from a mesenchymal-like pool of VSMC, though distinguishing between these phenotypes remains challenging^40^. Here we observed at D0 that a substantial amount of VSMC had already lost the contractile marker MYH11, and that many cells were expressing markers of fibroblasts, which is line with previous studies indicating that healthy veins are comprised of at least equal parts of VSMC and fibroblasts^41^. Our data also support the concept of initial VSMC remodeling and benign IH prior to vascular injury as part as the normal aging process^42^.

In line with the early presence of fibroblasts/myofibroblasts and activated cells, ECM remodeling emerged as another hallmark feature, with elevated expression of proteases (MMP9, Cathepsin K) and myofibroblast (HSP47, OPN, FN1), which facilitates ECM remodeling and IH progression^43^. MMPs, particularly MMP9, have previously been identified as critical drivers of pathological ECM remodeling in IH^44,45^.

Recent mouse studies also revealed osteochondrogenic transition of VSMCs in atherosclerosis^9,37,46^ and IH^24^. Here, we also observed evidence of VSMC reprogramming toward osteochondrogenic transition of VSMCs in human IH via the upregulation of key transcription factors (RunX2, Sox9)^47,48^. Osteochondrogenic differentiation is clinically relevant, contributing to vascular calcification, vessel stiffness, and reduced graft patency^25^. Thus, targeting osteochondrogenic pathways might represent a viable therapeutic approach to prevent vascular remodeling.

Our model further identified pronounced inflammatory responses as central feature of IH, with upregulation of pro-inflammatory cytokines and chemokines (IL-6, IL-1α, IL-1β, CCL5, CXCL2), which is a known feature of IH^5^. Of note, mouse studies in atherosclerosis models identified macrophage-like VSMC in atherosclerotic lesions contributing to the progression of atherosclerosis^38,46,49,50^. Here, the pro-inflammatory signature was mediated by activated VSMCs, EC, and fibroblasts but there was no evidence of further differentiation in macrophage-like cells expressing CD68, CD11b, or CD45^51,52^, suggesting that VSMC do not undergo macrophage dedifferentiation in the context of IH.

Oxidative and ER stress responses observed in our model provide mechanistic insights into IH progression. Elevated anti-oxidant proteins (HO-1, Nrf2, 4HNE) and ER stress indicators (XBP1, CHOP, ATF4) probably reflect cellular attempts to mitigate injury-induced damage. These findings are consistent with previous vascular injury studies showing that oxidative and ER stress responses drive VSMC apoptosis^53–56^. Interestingly, pre-clinical studies suggested that the ER-stress response promotes the phenotypic switch of VSMC towards the myofibroblast^57,58^.

Overall, our results emphasize the complex interplay between endothelial dysfunction, inflammation, oxidative stress, ECM remodeling, and cellular plasticity in IH pathology. Spatial transcriptomics effectively delineated specific transcriptional signatures, enhancing our understanding of localized cellular interactions and remodeling events.

## Limitations

Although this study provides a comprehensive characterization of molecular and cellular processes involved in IH formation using human vein segments, several limitations should be considered. First, the static *ex vivo* culture conditions employed lack physiological factors such as blood flow, pressure, and shear stress, all known to significantly influence IH development and vascular adaptation *in vivo*. Second, the absence of systemic immune components in the model limits our understanding of immune-mediated interactions and inflammation that play critical roles in saphenous vein graft disease pathogenesis. Third, due to limited sample size and examination at only two time points (Day 0 and Day 7), we could not fully capture the dynamic cellular and molecular processes occurring over an extended time course or correlate these changes with clinical outcomes. Future studies should aim to integrate dynamic flow conditions and explore immune-mediated mechanisms further.

In conclusion, our detailed characterization of human saphenous vein IH *ex vivo* highlights critical molecular and cellular processes underlying vein graft disease. This translationally relevant model provides valuable insights into the pathology and identifies novel therapeutic targets, particularly in VSMC plasticity and osteochondrogenic differentiation pathways, potentially improving clinical outcomes for vascular graft patients.

## a) Acknowledgments, Sources of Funding, & Disclosures

Acknowledgments: We are grateful to the Mouse Pathology Facility (MPF), the Cellular Imaging Facility (CIF), the Genomic Technologies Facility (GTF) and the Immune Landscape Laboratory (ILL) of the University of Lausanne for their support and expertise.

## Supplemental Material

Figure S1-S2

Major Resources Table

Tables S2–S3

Data Set1

## Non-standard Abbreviations and Acronyms

5hmC: 5-hydroxymethylcytosine
IH: Intimal hyperplasia
NI: neointima
M: media
L: lumen
DSP: Digital Spatial Profiler
VSMC: vascular smooth muscle cells
EC: endothelial cells
ECM: extracellular matrix
eNOS: endothelial nitric oxide synthase
VGEL: Verhoeff–van Gieson Elastic Lamina
FFPE: Formalin-Fixed Paraffin-Embedded
SV: saphenous vein
UPR: unfolded protein response

